# Super-resolution imaging of potassium channels with genetically encoded EGFP

**DOI:** 10.1101/2023.10.13.561998

**Authors:** Isabelle M. Call, Julian L. Bois, Scott B. Hansen

## Abstract

The plasma membrane is a well-organized structure of lipids and proteins, segmented into lipid compartments under 200 nm in size. This specific spatial patterning is crucial for the function of proteins and necessitates super-resolution imaging for its elucidation. Here, we establish that the genetically encoded enhanced green fluorescent protein (EGFP), when combined with direct optical reconstruction microscopy (dSTORM), tracks shear- and cholesterol-induced nanoscopic patterning of potassium channels overexpressed in HEK293T cells. Leveraging EGFP in dSTORM (EGFP-STORM), our findings indicate that cholesterol directs the C-terminus of TWIK-related potassium channel (TREK-1) to ceramide-enriched lipid ganglioside (GM1) clusters. In the absence of the C-terminus, the channel associates with phosphatidylinositol 4,5-bisphosphate (PIP_2_) cluster. Similarly, cholesterol derived from astrocytes repositions EGFP-tagged inward-rectifying potassium (Kir) channels into GM1 clusters. Without cholesterol, the channel aligns with PIP_2_ lipids. We deduce that cholesterol’s interaction with Kir sequesters the channel, separating it from its activating lipid PIP_2_. Fundamentally, a genetically encoded EGFP tag should make any protein amenable to dSTORM analysis.

## INTRODUCTION

Super-resolution imaging is a technique that employs single molecule localization (SML) to enhance the resolution of optical microscopy beyond the diffraction limit of light, which is approximately 250 nm. For this technique to be effective, a fluorophore must enter dark state with no fluorescence referred to as “blinking”. In 2008, it was discovered that this blinking property existed in common fluorophores, such as Cy3b, leading to the development of an SML adaptation to stochastic reconstruction microscopy (STORM) termed direct STORM (dSTORM) ^1^. dSTORM significantly streamlined the reagents required for SML since organic dyes suitable for this purpose were already accessible and frequently conjugated to commercially available antibodies.

Recently, dSTORM has been instrumental in discerning the association of proteins with distinct lipid compartments including, for example, in exploring membrane-mediated mechanisms underlying general anesthesia and mechanosensation. The anesthetic and mechanosensitive ion channel, TWIK-related potassium channel-1 (TREK-1), associates with ordered clusters of saturated lipids capable of binding palmitate – a 16-carbon saturated lipid covalently attached to proteins. Notably, anesthetics compete with palmitate at the anesthetic/palmitate (AP) site, as detailed in supplemental Figure 1^2^.

Many contemporary biological imaging techniques employ genetically encoded fluorophores, especially the enhanced green fluorescent protein (EGFP). EGFP is ideal for both live and fixed imaging as it negates the need for membrane permeabilization and antibodies, allowing access to intracellular molecules (as seen in Fig. 1A). Early developments led to the creation of a photoactivatable variant of EGFP, subsequently incorporated into a SML method termed photo-activated localization microscopy (PALM)^3^. Although these genetically encoded proteins excel in SML, they remain rarer than EGFP, and preparing samples for super-resolution imaging using them demands significant effort. Furthermore, recent advancements have adapted PALM techniques for dSTORM buffers, allowing the integration of photoactivatable encoded proteins with organic dyes. This development has been pivotal, as we’ve recently utilized dSTORM buffers in live cell imaging^4^.

**Figure 1.**
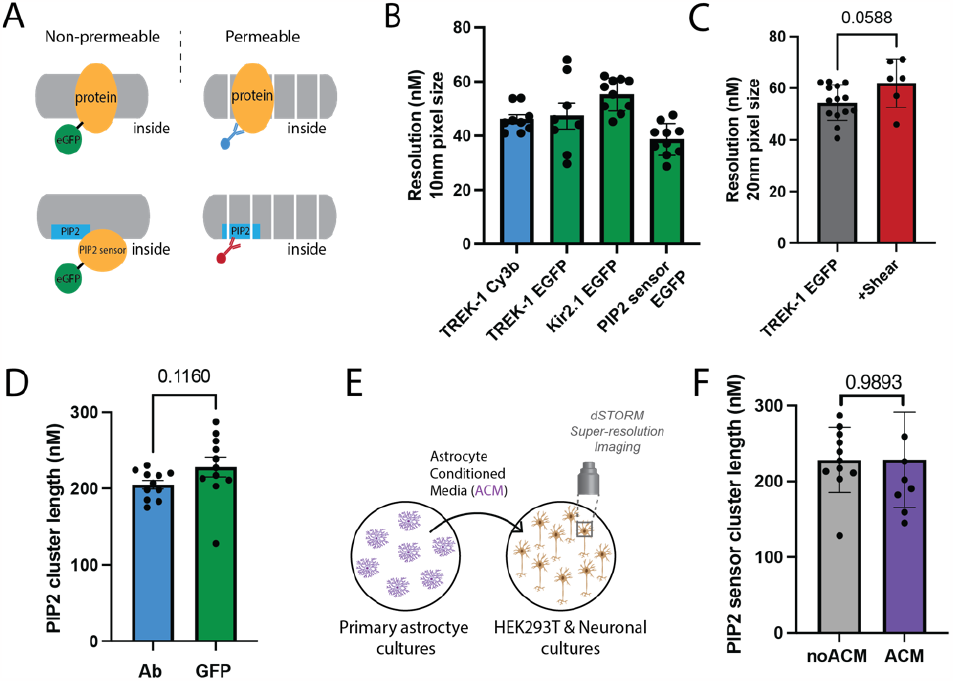
Characterization of genetically encoded EGFP as a dSTORM Fluorophore. (**A**) (Top) Cartoon depicting the differences in labeling techniques between labeling with enhanced green fluorescent protein (EGFP, green, left) and protein antibodies (Ab, blue, right) for direct stochastic reconstruction microscopy (dSTORM). Antibody labeling can require permeabilization (depicted with white lines). (Bottom) Experimental setup comparing a genetically encoded PIP_2_ sensor with EGFP (left) and anti-PIP_2_ antibody (right). The antibody typically requires permeabilization. (**B**) Ion channels TWIK-related potassium channel 1 (TREK-1) and inward rectifying potassium channel 2.1 (Kir_2.1_) were overexpressed in HEK293T cells with a genetically encoded EGFP tag. The resolution from dSTORM using EGFP tagged protein (green) is compared to the resolution using traditional Cy3B antibody-labeled TREK-1 (TREK-1_Cy3b, blue). When directly compared in side-by-side experiments, the resolutions were similar for both techniques (40-55 nm). (**C**) Fixation with 3 dynes/cm^2^ rotary shear marginally reduced the resolution, although the reduction was not statistically significant. (**D**) Cluster analysis of Atto647-conjugated anti-PIP_2_ antibody (Ab) and a EGFP tagged PIP_2_ sensor. (**E**) Cartoon depicting the experimental setup for treating a culture with astrocytes conditioned media (ACM). First ACM is generated by placing media on primary cortical astrocytes (purple cells, left) from mouse. The ACM is then removed, centrifuged, and placed on the treated culture. The treated cells are then fixed and imaged by direct stochastic reconstruction microscopy (dSTORM). (**F**) Determination of PIP_2_ lipid cluster sizes using EGFP encoded PIP_2_ sensor. The cluster size of PIP_2_ in HEK293T cells was unaffected by a 2-hour treatment with astrocyte-conditioned media.

Merging EGFP with dSTORM presents a promising avenue for SML. In theory, EGFP can be a formidable tool for ascertaining the nanoscopic spatial patterning of a protein. Proteins typically segregate into compartments of cholesterol and ganglioside lipids (GM1) or charged unsaturated lipids like PIP_2_ and PIP_3_ (as illustrated in Fig. S1A)^4–7^. To visualize the spatial organization of proteins alongside lipids, these lipids must also be fluorescently labeled. However, lipids cannot be genetically tagged - or at least not directly. This limitation underscores the need to incorporate organic dyes characteristic of dSTORM. A super-resolution technique that’s compatible with EGFP could significantly broaden the uptake of super-resolution imaging, particularly for elucidating nanoscopic protein patterning in intact cells. In this work, we employ EGFP-tagged TREK-1 and inward rectifying potassium channel 2.1 (Kir2.1) to showcase nanoscopic patterning alterations in response to fluid shear and increased membrane cholesterol levels.

## RESULTS

To demonstrate the capability of the genetically encoded EGFP for dSTORM, we tagged TREK-1 and the inward rectifying potassium channel 2.1 (Kir2.1) at their C-terminus with EGFP and overexpressed these proteins in HEK293T cells. Figure 1B presents a comparative analysis of the resolution achieved using dSTORM with EGFP-tagged proteins against the resolution from the traditionally labeled TREK-1 using a Cy3B antibody (referred to as TREK-1_Cy3b). When examined under identical conditions, both methodologies offered comparable resolutions, ranging from 40 to 55 nm for all tested proteins. This highlights the potential of EGFP as a reliable substitute for super-resolution imaging.

Notably, dSTORM excels in revealing shear-induced shifts in spatial patterning^2,4,8–11^. We discovered that EGFP provides a convenient means of quantifying shear-induced alterations in lipid localization. Introducing a 3 dynes/cm^2^ rotary shear during the fixation phase resulted in only a negligible, statistically non-significant drop in the resolution achieved through EGFP (refer to Fig. 1C). This supports the durability of EGFP across different experimental setups.

Furthermore, we employed a PIP_2_ sensor to measure the diameter of PIP_2_ clusters in HEK293T cells, and subsequently compared these measurements to sizes determined using a PIP_2_ antibody. This PIP_2_ sensor consists of the pleckstrin homology (PH) domain from delta phospholipase C tagged with EGFP. Cluster analysis using either the anti-PIP_2_ Ab or EGFP showed PIP_2_ cluster diameters of 205±6 nm and 228±42 nm, respectively (see Fig. 1D). These results suggest that the multivalent clustering of the PIP_2_ antibody does not alter the cluster diameter.

### EGFP dSTORM of Truncated TREK-1

The C-terminus of TREK-1 plays a pivotal role in many of the channel’s functional attributes. Notably, TREK-1 attaches to GM1 lipids via its C-terminus and forms a complex with phospholipase D2 (PLD2)^12^. This palmitoylated enzyme, PLD2, adheres to GM1 lipids and redirects TREK-1 away from PIP_2_^13^. Cholesterol amplifies PLD2’s interaction with GM1 lipids^4^, which consequently inhibits the channel. Notably, in the brain, cholesterol is supplied by astrocytes and transported to neurons to modulate protein function via GM1 lipids^14^.

The C-terminus of TREK-1 is also the target for most commercial antibodies. Excising the C-terminus for functional examination removes the epitope, thereby complicating dSTORM analysis. To circumvent this challenge, we opted for EGFP as the dSTORM fluorophore and overexpressed both the full length (TREK-FL) and a truncated yet functionally active version of TREK-1 (TREKtrunc) in HEK293T cells.

Figure 2A outlines the experimental protocol for the 3-color dSTORM used in our investigation. ACM, as described in Figure 1E, was introduced to cultured cells. Following fixation, permeabilization, and staining for GM1 and PIP_2_ clusters with CTxB and anti-PIP_2_ antibody respectively, we gauged the proximity of the two fluorophores using a pair correlation function^5^. Molecules in close proximity yielded a curve with a high pair correlation (as shown in Fig. 1B). The closest obtainable distance containing significant data (typically spanning 5-25 nm) was then compared.

**Figure 2:**
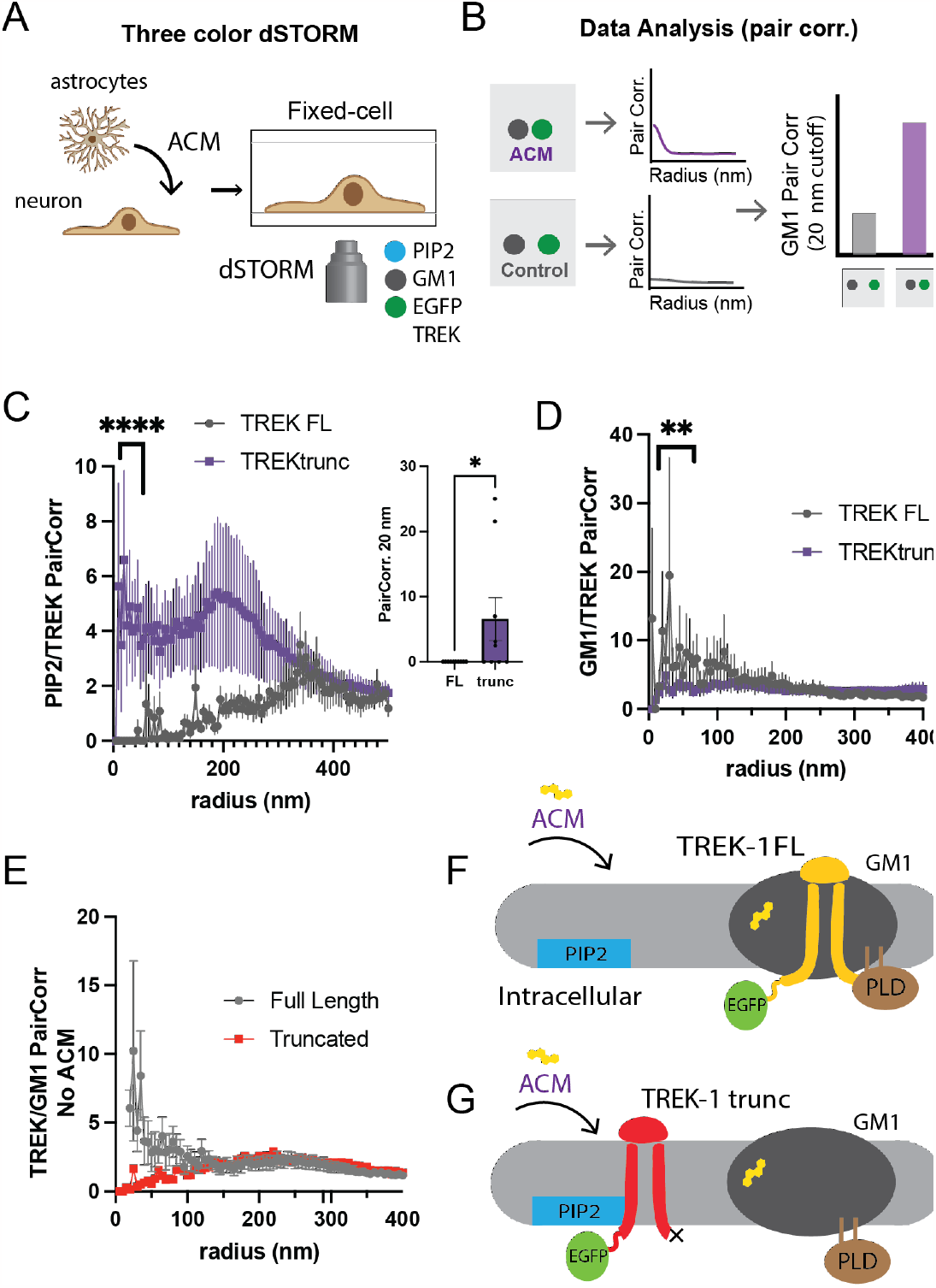
Tracking TREK-1 spatial patterning using EGFP-STORM. (**A**) An illustration demonstrates three-color direct stochastic optical reconstruction microscopy (dSTORM) technique employed in this study to determine the spatial patterning of human TWIK-related potassium channels (TREK-1) into lipid domains at nanoscopic distances (<250 nm). The proteins are genetically encoded with a C-terminal enhanced green fluorescent protein (EGFP) and over expressed in HEK293T cells. The cells are fixed, and the lipids are stained with fluorescent anti-PIP_2_ antibody (atto-647) and fluorescent cholesterol toxin B (cy3b-555). (**B**) The localization of a protein with a lipid is determined by pairwise correlation analysis (Pair corr.) of the GFP with each lipid (see hypothetical curve). A statistical comparison between two treatments is made at the shortest radius with sufficient data (see hypothetical bar graph). (**C-D**) Three-color dSTORM images of both the full-length TREK-1 (FL) and its C-terminally truncated variant (TREKtrunc). After treatments both with and without astrocyte-conditioned media (ACM), a cholesterol source in the brain, the cells were fixed and stained as shown in panel A. Panel C shows TREKtrunc maintained high pair correlation with PIP_2_ after ACM treatment (purple line), but not TREK FL. The inset shows statistical significance (n=9) at a single radius of 20 nm. Panel D shows that after ACM treatment, TREK FL correlated with GM1 lipids, but not TREKtrunc. (**E**) GM1 TREK-1 Pair corr. prior to cholesterol loading. In overexpressed cells TREK-1 associates with GM1 lipids in moderate cholesterol. (**F-G**) A summary cartoon illustrates the spatial patterning of TREK-1 using an EGFP tag. PIP_2_ clusters (blue) are shown separated from GM1 clusters (dark gray). Panel F shows when the concentration of membrane cholesterol (yellow) increases, the GFP signal associated with TREK-1 correlates with the GM1 lipids, but not PIP_2_. Panel G shows when TREK-1 is truncated, prior to its phospholipase d2 (PLD2) binding site (TREKtrunc, red shading), the channel associates strongly with PIP_2_, but not GM1 lipids.

Figure 2C juxtaposes TREKtrunc with TREK-FL. In the context of high cholesterol (i.e., ACM), TREK-1 exhibited a robust association with PIP_2_ (as seen in Fig. 2C) but not with GM1 lipids (Fig. 2D). Statistical assessment of _PIP2_/TREK-1 at 20 nm revealed a significant upsurge in the PIP_2_/TREKtrunc pair correlation as opposed to Wt. Conversely, TREK-FL displayed the opposite trend.

### Monitoring shear-induced changes in spatial patterning with EGFP

Previously, we demonstrated that shear forces prompt TREK-1 to shift from GM1 clusters to PIP_2_ clusters in permeabilized cells. However, due to the lack of a suitable antibody for the extracellular domain, direct comparisons between permeabilized and non-permeabilized cells were elusive. While photoactivatable fluorescent proteins are typically employed for such investigations, we evaluated this measurement using an EGFP tag and compared our findings with those from permeabilized cells.

Figure 3A presents TREK-1 under both shear and non-shear conditions in non-permeabilized cells. Notably, shear application significantly diminished the association of TREK-1 with GM1 clusters. For comparison, a similar experiment employing anti-TREK Cy3b in permeabilized cells is depicted in Figure 3C. A mouse phospholipase D2 (mPLD2), an enzyme known to activate TREK-1, showed a similar decrease in TREK-1 pair correlation with GM1 clusters upon shear application in HEK293T cells co-expressing TREK-1 and mPLD2.

**Figure 3.**
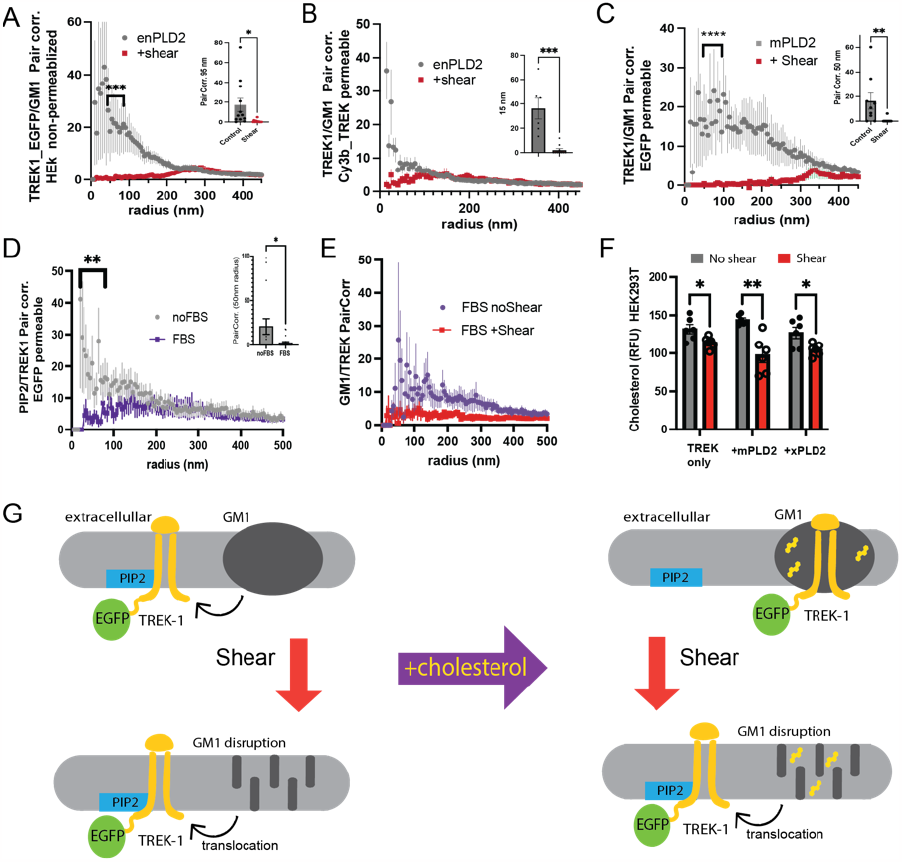
The use of EGFP to track shear-induced changes in Spatial Patterning of TREK-1. This figure demonstrates how an EGFP tag is used to monitor the trafficking of full-length human TREK-1 between PIP_2_ and GM1 lipids in response to shear and cholesterol in HEK293T cells. (**A**) Pairwise correlation (Pair corr.) of TREK_EGFP with GM1 lipids in non-permeabilized conditions. Prior to a 3 dynes/cm^2 fluid shear (red curve), there’s a correlation between overexpressed TREK-1 and GM1 lipids, which disappears post-shear. (**B**) An analogous experiment to (A), but employing cy3b-antiTREK-1 antibodies (cy3b_TREK) instead of EGFP encoding. (**C**) A variation of (B) where TREK-1 was co-expressed with mouse PLD2 (mPLD2), an activator of TREK-1. (**D-E**) Correlation analyses of EGFP-labeled TREK-1 in the presence of Fetal Bovine Serum (FBS, 10%) and apolipoprotein E (apoE, 4 ng/mL). (D) Reveals decreased TREK-EGFP correlation with PIP_2_ lipids in FBS (purple curve), hinting at TREK-1’s migration from PIP_2_ to GM1 clusters. (E) Confirms the GM1-TREK-1 correlation in FBS-treated cells. However, upon shear stress, this association breaks despite FBS presence, underscoring EGFP’s reliability in tracking TREK-1 spatial dynamics. (**F**) Cholesterol quantification via a cholesterol oxidase fluorescence assay shows a decline in cholesterol content in HEK293T cells post mechanical shear. (**G**) A schematic depicting TREK-1’s shift from GM1 (top) to PIP_2_ lipids (bottom) following mechanical shear (denoted by a red arrow). It also visualizes cholesterol’s role in TREK-1’s placement, either its removal through delipidated apoE (top left) or its addition via FBS (top right) in the absence of shear.

In line with the shear tests, EGFP proved effective for gauging cholesterol-induced relocations of TREK-1. Figure 3D reveals that cholesterol reduces PIP_2_/TREK-1 correlation while amplifying GM1/TREK-1 correlation. Intriguingly, shear application counteracted the effects of elevated cholesterol, as evidenced by EGFP-dSTORM. A subsequent cholesterol assay on the sheared samples displayed a reduction in free cholesterol, suggesting that the pair correlation decrease observed in Figure 3E might be attributable to a cholesterol concentration reversal.

### Spatial patterning of Kir2.1 with EGFP and mOrange fluorescent proteins

Kir2.1, a member of the inwardly rectifying potassium channel family, is instrumental in upholding the resting membrane potential and the repolarization phase of cardiac action potentials. Dysfunctions in Kir2.1 have been linked to several cardiac diseases^15^.

Kir2.1 activation requires the signaling lipid phosphatidylinositol 4,5-bisphosphate (PIP_2_)^16^. PIP_2_ not only binds to but directly regulates the channel. Without PIP_2_, the channel remains inactive^17^. Conversely, the content of cell membrane cholesterol has an inhibitory effect on Kir2, either directly or indirectly^18–20^. The exact molecular mechanisms behind cholesterol’s regulatory role remain elusive.

We hypothesized Kir2.1 inhibition occurs due to cholesterol-induced spatial patterning. Specifically, we hypothesized Kir2.1 undergoes nanoscopic clustering with cholesterol-rich ganglioside (GM1) lipids, distinct from its activating lipid, PIP_2_. As mentioned PIP_2_ clusters are typically separated from GM1 clusters. If Kir2.1 is associated to GM1 domains, it would be distanced from its agonist PIP_2_, thereby remaining inactive.

To test our hypothesis, we over expressed EGFP-Kir2.1 in HEK293T cells both with and without ACM. Using 3-color dSTORM imaging in HEK293T cells, we observed Kir2.1 movement after incubation with astrocyte conditioned media (ACM). Elevated cholesterol levels notably increased Kir2.1’s correlation with GM1 clusters (Fig. 4B, purple curve) while decreasing its correlation with PIP_2_ clusters (Fig. 4A, purple curve). This suggests cholesterol loading prompts Kir2.1 to shift away from PIP_2_ and toward GM1 clusters. Tagging Kir2.1 with mOrange, also allowed for Kir2.1/PIP_2_ pair correlations analysis (Fig. 4C) using a PIP_2_ antibody. The genetically endoded mOrange strongly correlated with an atto 488 conjugated PIP_2_ antibody. In theory the longer wavelength of mOrange, should allow for the colocalization of Kir2.1 with PIP_2_ using the EGFP-PIP_2_ sensor characterized in Figure 1. The two sensors did colocalize but not nearly to the same degree as with PIP_2_ antibody applied after fixing the cells. The overexpressed PIP_2_ sensory likely competes for Kir2.1 for PIP_2_ displacing some of it from PIP_2_ clusters and decreasing the correlation with of the two proteins.

**Figure 4:**
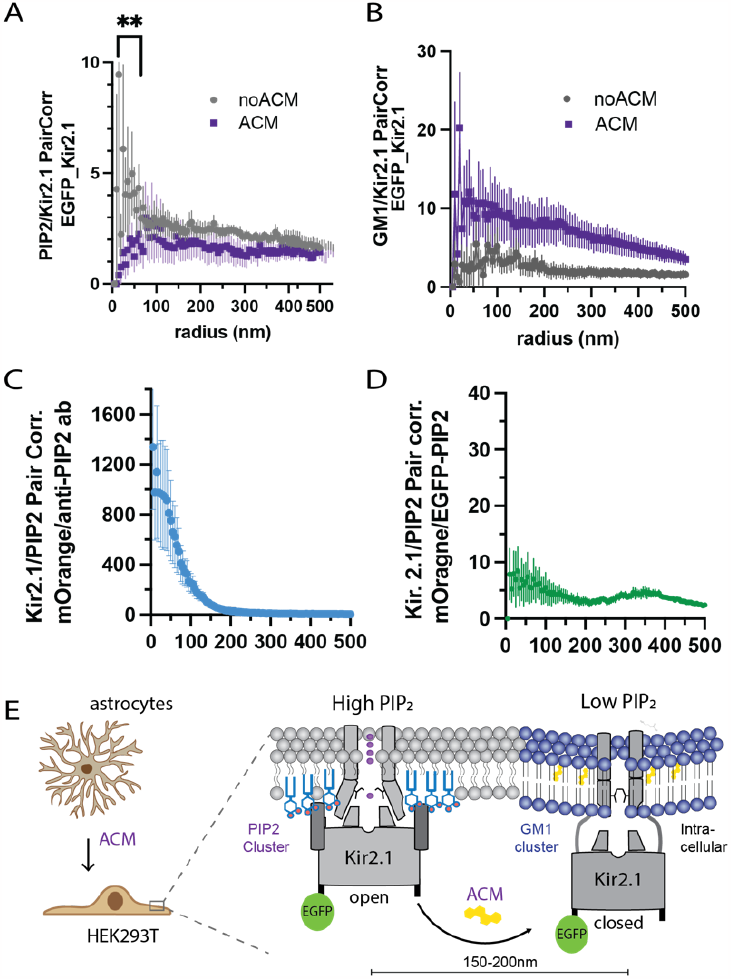
Effects of astrocyte cholesterol on inward rectifying K+ channel 2.1 (Kir2.1) spatial patterning. **(A)** Kir2.1’s pairwise correlation (PairCorr) with PIP_2_ was studied using dSTORM. Localization was assessed both in the presence and absence of astrocyte-conditioned media (ACM). Kir2.1 was tagged with EGFP, and the anti-PIP_2_ antibody was labeled with Atto647. **(B)** Correlation between GM1 and Kir2.1 extracted from the same samples as in (A). (**C**) Kir2.1’s correlation with PIP_2_, where Kir2.1 used an mOrange fluorescent tag and the anti-PIP2 antibody was conjugated with the atto-647 dye. (**D**) A proposed model illustrates Kir2.1’s cholesterol-driven inhibition due to its nanoscopic association with GM1 organized lipid clusters. The PIP2 clusters are set apart from GM1 clusters by a distance ranging from 150 to 200 nm. PIP_2_’s concentration is at its nadir within saturated GM1 clusters. Cholesterol induces TREK-1’s shift from the PIP_2_ cluster, aligning it with GM1 clusters instead. This localized dip in PIP_2_ concentration subsequently inhibits the channel.

## DISCUSSION

Our data collectively suggest that EGFP serves as an apt fluorophore for dSTORM in the context of shear and cholesterol-related experiments. The conclusions drawn from our shear experiments using EGFP are consistent with our previous findings. Furthermore, the technique’s applicability appears to be broad, as evidenced by the suitability of EGFP-Kir2.1 for dSTORM pair correlation analysis.

The most straightforward application is the two-color GM1/GFP pair correlation. Given that GM1 is extracellular, tracking a protein’s movement in and out of GM1 clusters with EGFP is likely to become a routine approach for palmitoylated proteins as well as other integral raft proteins. Our data show that EGFP-tagged TREK-1 yielded results comparable to those obtained using anti-TREK-1 antibodies.

Enhanced green fluorescent protein (EGFP) was engineered to augment the photostability of the original green fluorescent protein (GFP). EGFP boasts a quantum yield of 0.6 and an extinction coefficient of 55,000 M-1 cm-1. In contrast, the widely-used dye Alexa 647 (A647) has a quantum yield of 0.33 and an extinction coefficient of 239,000 M-1 cm-1. In addition to mOrange other fluorescent proteins likely have some utility in dSTORM.

Previous studies using PALM imaged PIP_2_domains by utilizing the mEOS-tagged PH domain. The measured cluster diameters in these studies were notably similar to our observations with EGFP^21^. However, minimal pair correlation was observed when the PIP_2_ sensor co-expressed with mOrange TREK-1. This diminished correlation might stem from competition between the sensor and TREK-1, suggesting a need for further experiments to adjust for expression levels.

In N2a cells, which naturally express TREK-1, we observed only a minor association with GM1 lipids. It’s noteworthy that TREK-1 is innately present in N2a cells.

With advancements in camera sensitivity, the requisite light intensity may decrease, potentially alleviating bleaching issues. In theory, this could pave the way for using EGFP in live cell imaging, further expanding the potential of dSTORM imaging techniques.

## METHODS

### Cell Culture and Gene Expression

HEK293T cells (ATCC Cat# CRL-3216, RRID:CVCL_0063) were maintained in DMEM (Corning cellgro) supplemented with 10% FBS, 100 units/mL penicillin, and 100 µg/mL streptomycin. For imaging experiments, cells were seeded onto poly-D-lysine-coated ibidi 8-well chamber microscope slides or 96-well plates. 24 hours post-seeding, transfections were performed using Turbofect (Thermo Scientific). The full-length human TREK-1 with a C-terminus GFP tag in a pCEH vector was a kind gift from Dr. Stephen Long. Mouse PLD2 constructs (mPLD2) and the inactive mutant (K758R, xPLD2) without a GFP tag in a pCGN vector were provided by Dr. Michael Frohman. For co-transfection experiments, TREK-1 was combined with mPLD2 or xPLD2 at a ratio of 1:4 (0.5µg of TREK-1 to 2µg of PLD DNA) ^12^.

### Antibody dye conjugation

Dye was reconstituted in 10 µL of DMSO. To prepare the dye-antibody mixture, 1.5 µL of the dye was combined with 50 µL of the respective antibody (TREK-1, Santa Cruz #sc-398449; PIP_2_, Echelon Biosciences #z-P045) and 6 µL of 1M NaHCO3 (pH 8.3). The mixture was incubated for 1.5 hours at room temperature in the dark. Post-incubation, a NAP-5 column was equilibrated with PBS. The antibody reaction was adjusted to a final volume of 200 µL with 140 µL PBS (pH 7.2) and loaded onto the NAP-5 column. After full absorption, the column was washed with 550 µL of PBS. The eluted fraction was collected post-wash and used for cell staining.

### Fixed cell preparation

80% confluent HEK293T cells were fixed with 3% paraformaldehyde and 0.1% glutaraldehyde for 10 minutes. Residual glutaraldehyde was neutralized using 0.1% NaBH4 for 7 minutes. Cells were washed thrice with PBS for 10 minutes per wash, followed by permeabilization using 0.2% Triton X-100 for 15 minutes. Blocking was done with 10% BSA/0.05% Triton/PBS at room temperature for 90 minutes. Fluorescent primary antibodies were added at a 1:100 dilution in 5% BSA/0.05% Triton/PBS and incubated for 60 minutes at room temperature. This was followed by five washes with 1% BSA/0.05% Triton/PBS for 15 minutes each. Subsequently, cells were post-fixed using the same fixing mixture for 10 minutes without agitation. Cells were then washed thrice with PBS for 5 minutes per wash and twice with dH2O for 3 minutes per wash.

### ACM treatment of cells

Primary mouse astrocytes from the cortex were cultivated in 10 cm plates using DMEM supplemented with 10% FBS. Upon reaching confluence, cells were sub-cultured. The spent media, termed astrocyte-conditioned media (ACM), was harvested every 3-4 days. Neuronal cells were treated with ACM for 1-2 hours prior to any additional procedures.

### Three-color Super-resolution dSTORM

Imaging was conducted on a Vutara VXL super-resolution microscope (Bruker Nano Surfaces, Salt Lake City, UT) that employs the 3D Biplane technique. The imaging setup utilized a Hamamatsu Orca Fusion BT sCMOS camera, paired with a 60x water-immersion objective (NA 1.2). Data analysis was performed using Vutara SRX software (version 7.0.07). Particles were localized in 3D using the software, which used a model function derived from previously recorded bead data sets. Imaging conditions included the use of 488, 561, and 647 nm lasers in a photoswitching buffer containing cysteamine, betamercaptoethanol, glucose oxidase (GLOX), catalase, Tris, and NaCl. The pH was maintained at 8.0. Direct antibody-dye conjugation was employed to avoid the need for fluorescent secondary antibodies. Image analysis was performed using the Statistical Analysis package in the Vutara SRX software (v7.0.07).

The pair correlation function g(r), cluster analysis, resolution analysis were performed using the Statistical Analysis package in the Vutara SRX software (v7.0.07). Pair correlation analysis is a statistical method used to determine the strength of correlation between two objects by counting the number of points of probe 2 within a certain donut-radius of each point of probe 1. This allows for localization to be determined without overlapping pixels as done in traditional diffraction-limited microscopy. For three-color EGFP-STORM probes 1 and 3 and 2 and 3 were also compared using the pair correlation function. Localization at super resolution is beyond techniques appropriate for diffraction-limited microscopy such as Pearson’s correlation coefficient. Lipid cluster size was determined using the DBSCAN clustering algorithm also included as part of the Vutara SRX software. And the resolution measurements were calculated by in the Vutara SRX software using resolution analysis with a 10 nm radius and smoothing.

### Cholesterol Oxidase protein preparation

The cholesterol oxidase gene from *Streptomyces sp*. (accession number AAA26719.1) was codon optimized and synthesized with a C-terminal 6x his tag (….KQDVTASHHHHH) and commercially cloned into a pET-28a(+) expression vector (GeneScript, USA). The plasmid was expanded in DH5*α* (kanamycin resistance) and used to transform BL21 cells by the heat shock method (note: very few colonies were observed). 500 mL starter culture in LB was grown and used to inoculate 3x1L of yeast tryptone media (YT) media pH 7. Protein expression was induced at an optical density (600 nm) of 0.6 with IPTG (Isopropyl β-D-1-thiogalactopyranoside) of 1mM. After 24 hours at room temperature, the cells were centrifuged (45 min 4500g), and resuspended in approximately 25 mL of lysis buffer (20 mM Tris-HCl pH 7.0 500mM NaCl). The cells were lysed by sonication, centrifuged, and the lysate, without protease inhibitors, was bound to a 1 mL cobalt talon metal affinity column by slow gravity flow over night at 4 °C. The next morning the column was washed with 10 mL of 20 mM imidazole in lysate buffer and then eluted with a gradient of 20 to 500 mM imidazole in the same buffer using an Acta Purifier. Approximately 30 mL corresponding to the cholesterol oxidase peak was concentration to 6.0 mg/mL and run on a Superdex 200 size exclusion column in 20 mM tris pH 7.0 to remove the imidazole.

### Cholesterol Assay

HEK293T cells were cultured in 48 well plates with 200uL media in each well and then changed to 200uL PBS for the shear treatment. The shear plate was incubated with PBS on an orbital rotator at 3dyn/cm^2 for 10 min in a 37°C incubator. The control plate was incubated with PBS for 10 min in the same incubator with no shear. Then the shear plate was incubated with 200uL 4%PFA+0.1% glutaraldehyde in PBS for 10 min with 3dyn/cm^2 shear and 10 min without shear. The control plate was fixed for 20 min with no shear.

### Statistical analysis

All statistical analysis were performed in GraphPad Prism 9.0. For the Student’s *t* test, significance was calculated using a two-tailed, unpaired parametric test with significance defined as \**P* < 0.05, \*\**P* < 0.01, \*\*\**P* < 0.001, and \*\*\*\**P* < 0.0001.

## Acknowledgments

We thank Michael Frohman from Stony Brook for the mouse PLD and mutant PLD cDNA and Steven Long from Memorial Sloan Kettering for human TREK-1-GFP. This work was supported by an R01 (R01NS112534) from the National Institutes of Health. We are grateful to the JPB Foundation for the purchase of a super resolution microscope. The authors declare no conflict of interest.

## Author Contributions

I.M.C. performed the dSTORM imaging, expressed and purified the cholesterol oxidase, and wrote the initial draft of the paper. J.L.B. performed cholesterol assays, wide field imaging experiments, and cluster analysis. S.B.H. designed the experiments, over saw the project, edited the draft, and assisted with data analysis.

## Supplemental figures

**Figure S1.**
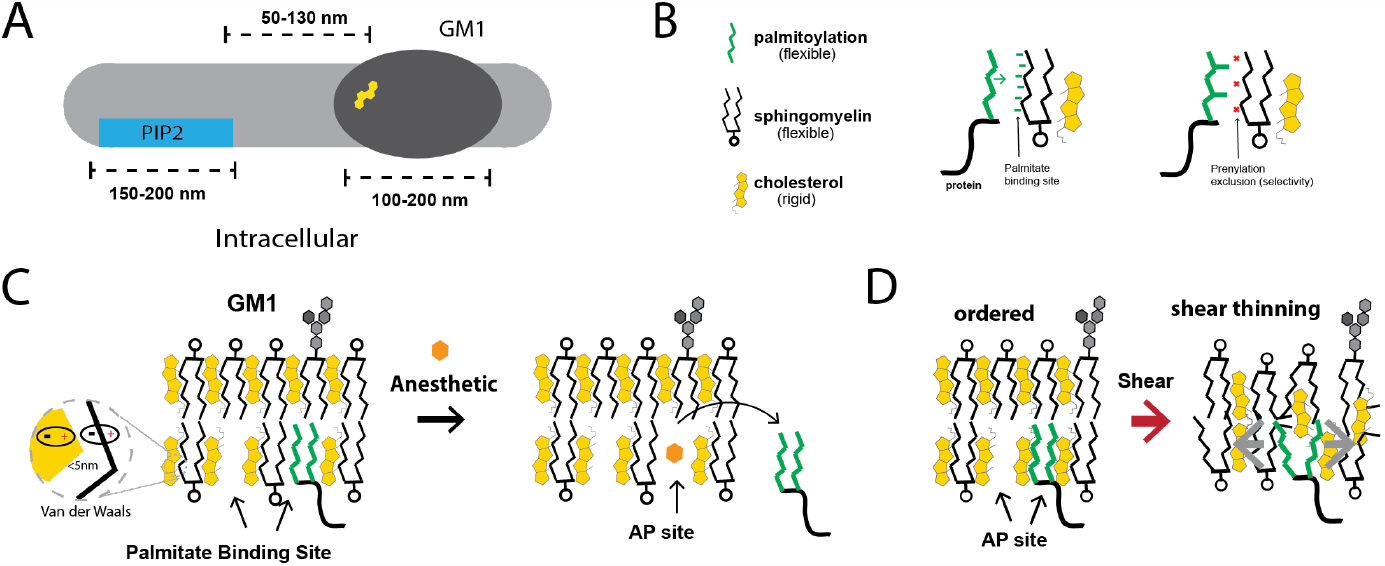
Structure and functional dynamics of glycolipid clusters. (**A**) Illustration showing the differentiation of glycolipid clusters comprised of phosphatidylinositol 4,5-bisphosphate (PIP_2_, blue, 150-200 nm in diameter) and the prototypical ganglioside monosialotetrahexosylganglioside (GM1, dark grey, 90-200 nm) in the plasma membrane. PIP_2_ lipid clusters emerge from charged protein interactions, while GM1 lipid clusters spontaneously form due to optimized hydrophobic packing. Cholesterol (yellow shading) partners with GM1 lipids, promoting the increased order of GM1 clusters^5,7,22,23^. (**B**) Breakdown of GM1 lipid components. Palmitate (green bars), a 16-carbon saturated lipid, can attach covalently to a protein (black line). Sphingomyelin (SM) shares a choline head group with phosphatidylcholine and GM1 has a saturated lipid structure like SM, but features a multi-sugar residue headgroup (grey hexagons). Palmitate binds with extended saturated lipids via Van der Waals forces, while prenylation (right), an unsaturated branched lipid, doesn’t bind to saturated GM1 lipids. (**C**) Illustration highlighting a palmitate binding site within GM1 domains and the competition with anesthetics for this site, termed here as the anesthetic/palmitate (AP) site. It’s specific to palmitate over prenylated lipids (left) and is also a target for anesthetics (right). Van der Waals forces between cholesterol and saturated SM are depicted. This ordered structure offers a palmitate binding surface, imparting function to GM1 domains. (**D**) Depiction of a GM1 domain sensing mechanical force via shear thinning. “Shear thinning” is a rheological concept where a viscous fluid reduces its viscosity in reaction to mechanical deformation. The left shows the membrane’s organized packing. Upon experiencing fluid shear, this order deforms. This diminishes Van der Waals forces between cholesterol and lipids, and consequently, the palmitate’s interaction with the structured surface. This deformation grants the lipids a more fluid nature, epitomizing shear thinning (illustrated on the right).

